# Effects of straw returning with potassium fertilizer on the stem lodging resistance, grain quality and yield of spring maize (Zea mays L.)

**DOI:** 10.1101/2021.04.19.440424

**Authors:** Ya-fang Fan, Ju-lin Gao, Ji-ying Sun, Jian Liu, Zhi-jun Su, Shu-ping Hu, Zhi-gang Wang, Xiao-fang Yu

## Abstract

The effects of straw returning with potassium fertilizer on the stem lodging resistance, grain quality and yield of spring maize were investigated to provide a scientific basis for the rational utilization of Inner Mongolia spring maize straw and potassium fertilizer resources. This study utilized Xianyu 335 as the test material, and a split plot design was carried out in three ecological regions from eastern to western Inner Mongolia (Tumochuan Plain irrigation area, Hetao Plain irrigation area and Lingnan warm dry zone), with the straw returning method as the main plot and potassium fertilizer dosage as the subplot. The stem resistance index, grain quality and yield were systematically identified. Both application of potassium fertilizer and straw returning improved the resistance and yield indicators of spring maize. Straw returning increased the effectiveness of potassium fertilizer application on spring maize plant height, ear height, fresh weight of stems, brix of stems and stem puncture strength by 2.82%-5.22%, 3.11%-5.90%, 15.96%-19.78%, 4.35%-4.50% and 8.89%-14.82%, respectively. Straw returning increased the effectiveness of potassium fertilizer application on the spring maize grain protein content, spring maize grain crude fat content, maize yield and yield variation coefficient by 3.49%-6.50%, 2.09%-4.43%, 4.87%-12.50% and 5.07%-7.55%, respectively. Straw returning can be combined with reasonable application of potassium fertilizer to increase the effectiveness of potassium fertilizer and enhance lodging resistance. Along with increased maize yield, straw returning also improves grain quality and enhances yield stability, providing a theoretical basis for high-yield and stress-resistant cultivation of Inner Mongolia spring maize, which can be popularized and applied in the spring maize planting areas of Inner Mongolia.

## Introduction

As the most popular grain crop in China, the yield and quality of maize (Zea mays L.) must be maintained at high levels to ensure stable increases in national grain production and food security [1]. With the continuous adjustment and improvement of the goals and tasks of the maize planting industry in China, priorities have changed from the pursuit of high yield alone to a broader focus on optimizing structure and enhancing quality while controlling cost. Therefore, current agricultural strategies are aimed at adjusting planting modes, improving the quality of maize products, and promoting the ecologically responsible and high-quality development of the planting industry on the basis of ensuring a stable increase in grain production.

Lodging is a major factor that limits maize yields. According to reports from the previous decade [2,3], the lodging rate of maize is significantly positively correlated with ear height, and yield decreases by 108 kg/hm^2^ for every increase of 1% in the lodging rate during maize production. Mechanical properties associated with maize stem lodging resistance are important indicators of the degree of lodging resistance of maize plants and are significantly negatively correlated with field lodging rates. The phenotypic traits of stems influence mechanical properties associated with lodging resistance and thus determine the lodging resistance of plants [4-6]. In response to the direction in which grain production practices are developing in China, stable production and guaranteed income require enhancement of grain development, improvement in maize grain quality, and increased economic efficiency. The quality of maize grain is altered by long-term fertilization [7]. In addition, grain quality is influenced by genetics, fertilization measures and environmental conditions [8]. The practice of returning straw to maize fields provides abundant organic substances, which can promote the synthesis of crude fat, protein and starch in maize grain and improve grain quality [9].

The application of potassium fertilizer can improve maize resistance to lodging, enhancing quality and increasing maize yield [10]. However, due to a lack of potassium resources in China, potassium fertilizer is generally imported, leading to high cost and limited availability. The effectiveness of K^+^ in soil and fertilizer is dependent on the soil texture [11]. To overcome the increasingly serious problem of soil potassium deficiency, effective means of supplementing soil potassium in addition to fertilizer potassium are required. With the current level of soil productivity, there is a need to explore the production potential of the soil itself in order to increase the nutrient level of the soil, improve the soil structure and physicochemical properties, optimize the ecological environment of the farmland, and maintain high crop yields, with the goal of avoiding resource waste and environmental pollution [12-14]. The direct burning, abandonment or incineration of straw resources causes significant environmental pollution and represents a significant waste of resources. Therefore, straw returning is a method of achieving comprehensive utilization of straw resources [15]. Straw returning can optimize soil structure and physical and chemical properties, improve soil enzyme activity, and increase soil nutrients and maize yield [16,17]. China is rich in straw resources. The average annual yield of maize straw in China is 399.18 million t, and the potassium nutrient content provided by maize straw returning in China is 4.79 million t K_2_O. The potassium fertilizer substitution potential of maize straw returning in the growing season in China is 24.4 kg/hm^2^ K_2_O [18], and the release rate of potassium during this season is approximately 85% [19]. It is generally believed that the use of straw resources can alleviate soil potassium deficiency and enrich the soil potassium pool, which should improve soil fertility and allow growers to meet the potassium demand of maize production in China.

Previous studies mostly focused on adding potassium fertilizer to increase the yield of maize based on straw returning [20,21]. However, there are few studies on the effects of straw returning with potassium fertilizer on the stem lodging resistance, grain quality and yield of spring maize. The effects of straw returning with potassium fertilizer on the phenotypic traits, stem lodging resistance mechanical properties, maize grain quality and yield of spring maize were investigated in three ecological regions from eastern to western Inner Mongolia, and a cultivation model suitable for high-quality and high-yield maize agriculture was explored. This study provides a basis for the cultivation of high-yield and stress-resistant spring maize in Inner Mongolia and elsewhere in China, as well as the development of a high-quality green planting industry.

## Materials and Methods

### Overview of the test site

Three ecological regions in Inner Mongolia (Tumochuan Plain irrigation area, Hetao Plain irrigation area and Lingnan warm dry zone) were used as test sites in 2019. The longitude and latitude, sunshine hours from April to October, average temperature, and rainfall at each test site are listed in Table 1. Table 2 shows the soil type and basic soil fertility of each test site.

**Table 1.** Latitude and longitude and climatic conditions of three ecological regions in Inner Mongolia

| Ecological region | Experimental sites | Latitude | Longitude | Solar radiation (hour) | Average temperature (°C) | Precipitation (mm) |
| --- | --- | --- | --- | --- | --- | --- |
| Hetao Plain irrigation area | Bayannur | 41°11'N | 122°49'E | 1924.3 | 20.4 | 157.6 |
| Tumochuan Plain irrigation area | Baotou | 40°32'N | 122°48'E | 1716.4 | 21.1 | 317.7 |
| Lingnan warm dry zone | Xing'an | 46°45'N | 122°47'E | 1330.5 | 22.3 | 392.5 |

**Table 2.** Soil type and soil basic fertility of three ecological regions in Inner Mongolia

| Ecological region | Soil type | Organic Matter (g/kg) | Total N (g/kg) | Available N (mg/kg) | Olsen P (mg/kg) | Available K (mg/kg) | pH |
| --- | --- | --- | --- | --- | --- | --- | --- |
| Hetao Plain irrigation area | Irrigated soil | 26 | 0.4 | 87.5 | 6.6 | 140.3 | 7.9 |
| Tumochuan Plain irrigation area | Silty loam | 26.7 | 0.5 | 92.4 | 8.8 | 118 | 7.6 |
| Lingnan warm dry zone | Black soil | 29.1 | 1.3 | 100.6 | 13.7 | 112.8 | 7.8 |

The field study was carried out on the official land which belonged to the key laboratory of crop cultivation and genetic improvement of Inner Mongolia Autonomous Region, permission was given after research application passing verification. During the field study none of endangered or protected species were involved. No specific permissions were required for conducting the field study because it was not carried out in protected area.

### Experimental design

This study utilized Xianyu 335 as the test material in a split plot design, with the straw returning method as the main plot and potassium fertilizer dosage as the subplot. The 4 treatments used in the study were straw returning + potassium fertilizer (ST+6K), straw returning + no potassium fertilizer (ST+0K), no straw returning + potassium fertilizer (NST+6K), and no straw returning + no potassium fertilizer (NST+0K). The row length was 30 m, the row width was 5 m, and the row spacing was 0.60 m. Five replicates were used in this experiment, and the planting density was 82500 plants/hm^2^. The straw returning treatments utilized pulverized straw that was returned to the field in the autumn of the previous year. The non-returning treatments were all household shallow rotation modes. Potassium was applied as 90 kg/hm^2^ potassium sulfate (K_2_O 50%) and 228 kg/hm^2^ diammonium phosphate (P_2_O_5_ 46%) once as the base fertilizer before sowing. For the treatments without potassium, only 228 kg/hm^2^ diammonium phosphate (P_2_O_5_ 46%) was applied once as the base fertilizer before sowing. The top dressing of each treatment was 652 kg/hm^2^ (N46%), which was applied in the jointing stage and the bell stage at a ratio of 3:7. Other management procedures followed typical field production practices.

### Measurement items and methods

Before sowing, 0-20 cm soil samples were taken for each treatment, ventilated and dried in a cool place, after which they were ground to pass through a 0.15-0.25 mm soil sieve. According to the measurement requirements, soil samples with different particle sizes were used to determine soil essential nutrients [22].

(1) Soil organic matter determination was performed using the potassium dichromate titration method. (2) Soil total nitrogen determination was performed using a Kjeldahl nitrogen analyzer (K-9840, Jinan) and the semi-micro Kjeldahl method. (3) Soil available phosphorus determination was performed using the NaH_2_CO_3_ (0.5 mol/L) Mo-Sb colorimetric method. (4) Soil available potassium determination was performed using the NH_4_Ac (1 mol/L) extraction 30-min flame photometric method. (5) Soil alkaline hydrolysis determination was performed using the alkaline hydrolysis diffusion-absorption method.

The following stem indicators were measured during the silking period. (1) Plant height was measured by using a steel ruler to measure the distance from the top of the tassel to the ridge side. (2) Ear height was measured by using a steel ruler to measure the distance between the first ear internode and the ridge side. (3) Stem diameter was measured by using a Vernier caliper to measure the third stem node at the stem base part. (4) Ear stem length was determined by measuring the maize ear node length. (5) Stem fresh weight was determined by measuring the maize stem fresh weight. (6) Stem dry weight was determined by measuring the weight of dried maize stems. (7) The water content of the stems was calculated as the ratio of (stem fresh weight–stem dry weight) and stem fresh weight. (8) To measure the brix of the stems, the maize stems were extracted and mixed, and 1-2 mL of the mixture was measured with a handheld digital sugar meter (PAL-1, Japan ATAGO, accuracy = ±0.2%). (9) To assess stem lodging resistance mechanical indicators, the stem puncture strength, compressive strength and bending strength of the third stem node at the maize stem base were measured with a plant stem strength instrument (YYD-1, Tuopu Yunnong, Zhejiang, accuracy = ±0.5% F.S.).

The starch content, crude fat content, protein content, and water content of maize grains were measured with a FOSS near-infrared grain quality analyzer (Infratee TM 1241, FOSS, Denmark) at maturity.

At the physiological maturity stage, two rows in the middle of the measured production area were selected, and all plants in these rows were harvested after removal of the side plants. The number of harvested ears was counted. Ten plants with uniform ear growth were selected for determination of ear rows, row grains, 1000-grain weight, and grain water content (measured with an LDS-1G moisture content detector), which were converted into maize yield (converted into hectare yield with 14% water content).

### Data statistical analysis

Data SPSS window version 17 (SPSS Inc., Chicago, USA) was used to finishing statistical analysis. Under straw-return treatments, potassium fertilizer treatments, and ecological regions, we examine stem lodging resistance, grain quality and yield of spring maize using GLM based on the model for a split-plot design [23]. The values were all the mean squares (MS) of the ANOVA. Straw-return treatments, potassium fertilizer treatments, and ecological regions were the independent variables, and the stem lodging resistance, grain quality and yield of spring maize were dependent variables in this test. In order to determine the impact of independent variables on dependent variables, statistically significant variance was tested using three-way analysis of variance, and multiple comparisons were made using the least significant difference (LSD) test with α = 0.05 [24]. Histograms were conducted by using Sigma Plot 12.5. And different letters on histograms indicated that means statistically different at P<0.05 level.

## Results

### Effects of straw returning combined with potassium fertilizer on the morphological indexes of spring maize stems

As shown in Table S1, the effects of variation of the straw returning method, potassium fertilizer dosage, and ecological region on plant height, ear height, stem diameter and ear stem length reached an extremely significant level. The effects of the interaction of the straw returning method and potassium fertilizer dosage on the above-mentioned indicators reached a very significant level. The effect of the interaction of the straw returning method and ecological region and the interaction of the potassium fertilizer dosage and ecological region on plant height, stem diameter and ear stem length reached a significant or extremely significant level. The effect of the interaction of the straw returning method, potassium fertilizer dosage and ecological region on plant height reached an extremely significant level.

**Table S1.** ANOVA results for maize stem morphological indicators under different straw returning methods and potassium fertilizer treatments

| Source of variation | Plant height<br>(cm) | Ear height<br>(cm) | Ear height<br>coefficient | Stem meter<br>(mm) | Stem length<br>(cm) |
| --- | --- | --- | --- | --- | --- |
| Straw returning method (S) | 1539.66** | 160.66** | ns | 19.80** | 10.53** |
| Potassium fertilizer dosage (K) | 6119.79** | 1060.42** | ns | 44.63** | 23.18** |
| Ecological region (E) | 444.16** | 153.68** | ** | 41.34** | 0.95** |
| S×K | 540.72** | 99.12** | ns | 6.19** | 1.08** |
| S×E | 67.48** | 8.01ns | ns | 0.16* | 0.18** |
| K×E | 21.30** | 2.86ns | ns | 0.34** | 0.03* |
| S×K×E | 19.92** | 4.08ns | ns | 0.04ns | ns |

Note: \* and \*\* represent the significance at the level of 5% and 1%, respectively, while ns represents an insignificant difference. “S, K and E” represent straw-return treatments, potassium fertilizer treatments and ecological regions, respectively. These values in the table are the mean squares (MS) of the ANOVA. The same below.

As shown in Table 3, maize plant height, ear height, ear stem diameter and ear stem length varied among the treatment groups. In the Tumochuan Plain irrigation area, under the straw returning treatment, the maize plant height, ear height, ear height coefficient, stem diameter and ear stem length increased by 7.74%, 8.36%, 0.57%, 8.57%, and 11.24%, respectively, with potassium application in comparison with no potassium application. With no straw returning treatment, the maize plant height, ear height, ear height coefficient, stem diameter and ear stem length increased by 4.92%, 5.25%, 0.31%, 4.14%, and 7.21%, respectively, with potassium application in comparison with no potassium application. Straw returning improved the effectiveness of potassium application on the maize plant height, ear height, ear height coefficient, stem diameter and ear stem length by 2.82%, 3.11%, 0.26%, 4.43% and 4.03%, respectively.

**Table 3.** Effects of the interaction of the straw returning method and potassium fertilizer dosage on the morphological indicators of maize stems in different ecological regions

| Ecological region | Straw returning method | Potassium fertilizer dosage | Plant height (cm) | Ear height(cm) | Ear height coefficient | Stem meter(mm) | Stem length(cm) |
| --- | --- | --- | --- | --- | --- | --- | --- |
| Tumochuan Plain<br>irrigation area | ST | 6K | 329.72±3.82a | 128.17±2.33a | 0.39±0.01a | 27.58±1.15a | 14.40±0.11a |
|  |  | 0K | 306.03±3.87c | 118.29±2.34c | 0.39±0.01a | 25.41±1.16c | 12.95±0.13c |
|  | NST | 6K | 318.10±4.69b | 124.43±2.43b | 0.39±0.01a | 26.01±0.85b | 13.16±0.10b |
|  |  | 0K | 303.17±3.73d | 118.23±3.01c | 0.39±0.01a | 24.98±0.91d | 12.28±0.24d |
|  | Under ST, the percentage increase of 6K compared to 0K % |  | 7.74 | 8.36 | 0.57 | 8.57 | 11.24 |
|  | Under NST, the percentage increase of 6K compared to 0K % |  | 4.92 | 5.25 | 0.31 | 4.14 | 7.21 |
|  | ST increases the percentage of K effectiveness % |  | 2.82 | 3.11 | 0.26 | 4.43 | 4.03 |
| Hetao Plain<br>irrigation area | ST | 6K | 340.63±2.28a | 133.59±2.71a | 0.39±0.01a | 29.75±0.84a | 14.63±0.13a |
|  |  | 0K | 309.84±2.63c | 120.73±2.13c | 0.39±0.01a | 27.12±0.96c | 13.06±0.14c |
|  | NST | 6K | 318.14±4.35b | 125.61±2.09b | 0.39±0.01a | 27.80±0.86b | 13.77±0.18b |
|  |  | 0K | 303.81±4.68d | 119.91±1.76c | 0.39±0.01a | 26.38±1.11d | 12.68±0.19d |
|  | Under ST, the percentage increase of 6K compared to 0K % |  | 9.94 | 10.65 | 0.64 | 9.74 | 12.03 |
|  | Under NST, the percentage increase of 6K compared to 0K % |  | 4.72 | 4.75 | 0.03 | 5.39 | 8.62 |
|  | ST increases the percentage of K effectiveness % |  | 5.22 | 5.90 | 0.62 | 4.35 | 3.41 |
| Lingnan warm dry<br>zone | ST | 6K | 325.24±2.57a | 126.27±3.02a | 0.39±0.01a | 26.61±1.05a | 14.35±0.11a |
|  |  | 0K | 301.11±3.81c | 116.08±2.75c | 0.39±0.01a | 24.32±0.95c | 12.84±0.13c |
|  | NST | 6K | 310.95±4.11b | 120.47±1.29b | 0.39±0.00a | 24.77±0.74b | 13.13±0.06b |
|  |  | 0K | 297.62±3.98d | 114.85±1.44c | 0.39±0.00a | 23.96±0.70d | 12.18±0.11d |
|  | Under ST, the percentage increase of 6K compared to 0K % |  | 8.02 | 8.78 | 0.71 | 9.42 | 11.76 |
|  | Under NST, the percentage increase of 6K compared to 0K % |  | 4.48 | 4.90 | 0.40 | 3.36 | 7.80 |
|  | ST increases the percentage of K effectiveness % |  | 3.54 | 3.88 | 0.31 | 6.07 | 3.96 |

Note: These values in the table are the mean squares (MS) of the ANOVA. Values followed by different letters in a column are significant among treatments at the 5% level. The same below.

In the Hetao Plain irrigation area, with the straw returning treatment, the maize plant height, ear height, ear height coefficient, stem diameter and ear stem length increased by 9.94%, 10.65%, 0.64%, 9.74%, and 12.03%, respectively, with potassium application in comparison with no potassium application. With no straw returning, the maize plant height, ear height, ear height coefficient, stem diameter and ear stem length increased by 4.72%, 4.75%, 0.03%, 5.39%, and 8.62%, respectively, with potassium application in comparison with no potassium application. Straw returning increased the effectiveness of potassium application on the maize plant height, ear height, ear height coefficient, stem diameter and ear stem length by 5.22%, 5.90%, 0.62%, 4.35% and 3.41%, respectively.

In the Lingnan warm dry zone, under the straw returning treatment, the maize plant height, ear height, ear height coefficient, stem diameter and ear stem length were increased by 8.02%, 8.78%, 0.71%, 9.42%, and 11.76%, respectively, with potassium application in comparison with no potassium application. With no straw returning, the maize plant height, ear height, ear height coefficient, stem diameter and ear stem length were increased by 4.48%, 4.90%, 0.40%, 3.36%, and 7.80%, respectively, with potassium application in comparison with no potassium application. Straw returning increased the effectiveness of potassium application on the maize plant height, ear height, ear height coefficient, stem diameter and ear stem length by 3.54%, 3.88%, 0.31%, 6.07% and 3.96%, respectively.

### Effects of straw returning with potassium fertilizer on the phenotypic traits of spring maize stems

As shown in Table S2, the effects of variation of the straw returning method, potassium fertilizer dosage, and ecological region on the stem fresh weight, stem dry weight, water content of maize grain and brix of stems reached an extremely significant level. The effects of the interaction of the straw returning method and potassium fertilizer dosage on the stem fresh weight, stem dry weight, water content of maize grain and brix of stems reached a significant or extremely significant level. The effects of the interaction of the straw returning method and ecological region and the interaction of the potassium fertilizer dosage and ecological region on the stem fresh weight, stem dry weight and water content of stems reached a significant or extremely significant level. The effect of the interaction of the straw returning method, potassium fertilizer dosage and ecological region on the stem fresh weight reached a significant level.

**Table S2.**
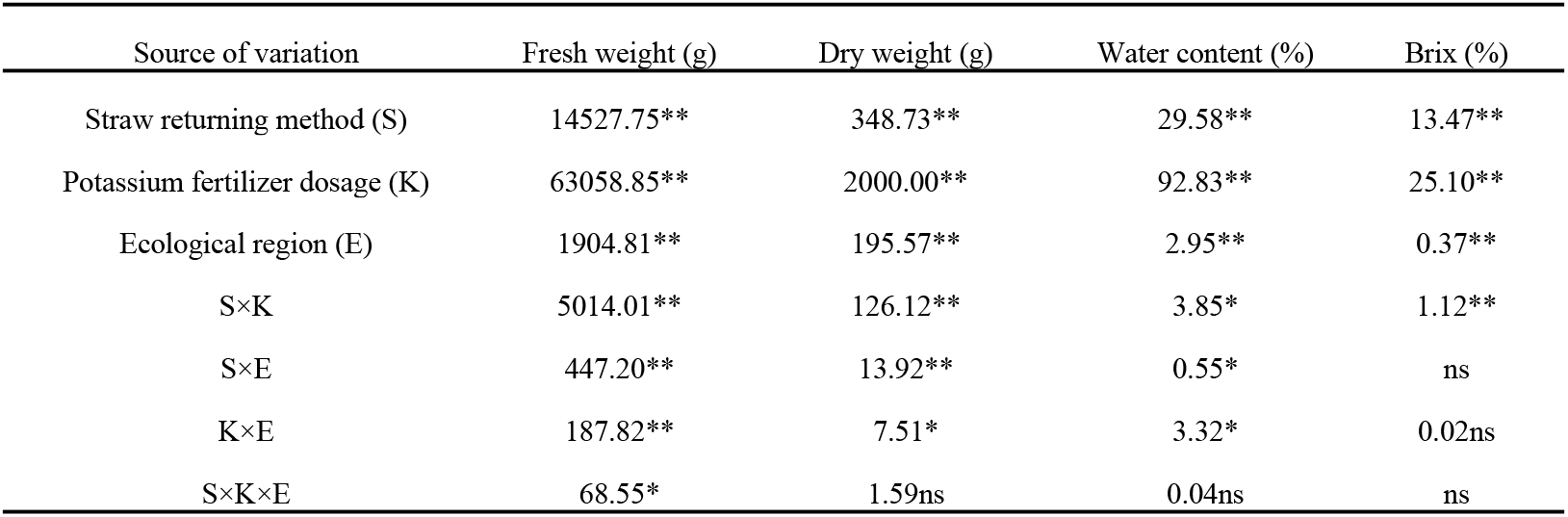
ANOVA results for maize stem phenotypic traits under different straw returning methods and potassium fertilizer treatments

| Source of variation | Fresh weight (g) | Dry weight (g) | Water content (%) | Brix (%) |
| --- | --- | --- | --- | --- |
| Straw returning method (S) | 14527.75** | 348.73** | 29.58** | 13.47** |
| Potassium fertilizer dosage (K) | 63058.85** | 2000.00** | 92.83** | 25.10** |
| Ecological region (E) | 1904.81** | 195.57** | 2.95** | 0.37** |
| S×K | 5014.01** | 126.12** | 3.85* | 1.12** |
| S×E | 447.20** | 13.92** | 0.55* | ns |
| K×E | 187.82** | 7.51* | 3.32* | 0.02ns |
| S×K×E | 68.55* | 1.59ns | 0.04ns | ns |

As shown in Table 4, the maize stem fresh weight, stem dry weight, water content of stems and brix of stems differed among the treatments. In the Tumochuan Plain irrigation area, under the straw returning treatment, the maize stem fresh weight, stem dry weight, water content of stems and brix of stems increased by 44.20%, 29.98%, 3.66%, and 15.32%, respectively, with potassium application in comparison with no potassium application. With no straw returning treatment, the maize stem fresh weight, stem dry weight, water content of stems and brix of stems increased by 28.22%, 21.09%, 2.11%, and 10.98%, respectively, with potassium application in comparison with no potassium application. Straw returning increased the effectiveness of potassium application on the maize stem fresh weight, dry weight, water content and brix of stems by 15.98%, 8.89%, 1.55% and 4.35%, respectively.

**Table 4.** Effects of the interaction of the straw returning method and potassium fertilizer dosage on the phenotypic traits of maize stems in different ecological regions

| Ecological region | Straw returning method | Potassium fertilizer dosage | Fresh weight (g) | Dry weight (g) | Water content (%) | Brix(%) |
| --- | --- | --- | --- | --- | --- | --- |
| Tumochuan Plain irrigation area | ST | 6K | 273.16±3.01a | 66.73±2.77a | 75.57±0.99a | 12.37±0.20a |
|  |  | 0K | 189.44±2.82c | 51.33±1.58c | 72.90±0.88c | 10.73±0.11c |
|  | NST | 6K | 230.17±3.67b | 60.23±2.18b | 73.82±1.31b | 11.15±0.26b |
|  |  | 0K | 179.52±1.59d | 49.74±1.86d | 72.29±0.98d | 10.05±0.12d |
|  | Under ST, the percentage increase of 6K compared to 0K % |  | 44.20 | 29.98 | 3.66 | 15.32 |
|  | Under NST, the percentage increase of 6K compared to 0K % |  | 28.22 | 21.09 | 2.11 | 10.98 |
|  | ST increases the percentage of K effectiveness % |  | 15.98 | 8.89 | 1.55 | 4.35 |
| Hetao Plain irrigation area | ST | 6K | 294.74±4.82a | 71.17±3.27a | 75.85±1.07a | 12.44±0.12a |
|  |  | 0K | 202.76±4.89c | 56.98±1.65c | 71.89±0.78c | 10.93±0.16c |
|  | NST | 6K | 230.20±7.43b | 60.89±2.11b | 73.53±1.07b | 11.25±0.06b |
|  |  | 0K | 183.29±3.06d | 53.78±1.70d | 70.64±1.28d | 10.29±0.11d |
|  | Under ST, the percentage increase of 6K compared to 0K % |  | 45.41 | 24.88 | 5.51 | 13.82 |
|  | Under NST, the percentage increase of 6K compared to 0K % |  | 25.63 | 13.23 | 4.10 | 9.38 |
|  | ST increases the percentage of K effectiveness % |  | 19.78 | 11.65 | 1.42 | 4.44 |
| Lingnan warm dry zone | ST | 6K | 257.51±3.52a | 63.21±2.11a | 75.44±1.10a | 12.22±0.13a |
|  |  | 0K | 183.85±3.76c | 49.45±1.64c | 73.09±1.12c | 10.67±0.17c |
|  | NST | 6K | 216.83±3.52b | 56.82±1.15b | 73.79±0.95b | 10.97±0.09b |
|  |  | 0K | 174.73±2.71d | 48.48±1.83d | 72.25±1.02d | 9.97±0.07d |
|  | Under ST, the percentage increase of 6K compared to 0K % |  | 40.09 | 27.82 | 3.22 | 14.51 |
|  | Under NST, the percentage increase of 6K compared to 0K % |  | 24.13 | 17.28 | 2.13 | 10.02 |
|  | ST increases the percentage of K effectiveness % |  | 15.96 | 10.55 | 1.09 | 4.50 |

In the Hetao Plain irrigation area, under the straw returning treatment, the maize stem fresh weight, stem dry weight, water content and brix of stems increased by 45.41%, 24.88%, 5.51%, and 13.82%, respectively, with potassium application in comparison with no potassium application. With no straw returning treatment, the maize stem fresh weight, stem dry weight, water content and brix of stems increased by 25.63%, 13.22%, 4.10%, and 9.38%, respectively, with potassium application in comparison with no potassium application. Straw returning increased the effectiveness of potassium application on the maize stem fresh weight, stem dry weight, water content and brix of stems by 19.78%, 11.65%, 1.42%, and 4.44%, respectively.

In the Lingnan warm dry zone, under the straw returning treatment, the maize stem fresh weight, stem dry weight, water content and brix of stems increased by 40.09%, 27.82%, 3.22%, and 14.51%, respectively, with potassium application in comparison with no potassium application. With no straw returning treatment, the maize stem fresh weight, stem dry weight, water content and brix of stems were increased by 24.13%, 17.28%, 2.13%, and 10.02%, respectively, with potassium application in comparison with no potassium application. Straw returning increased the effectiveness of potassium application on the maize stem fresh weight, stem dry weight, water content and brix of stems by 15.96%, 10.55%, 1.09%, and 4.50%, respectively.

### Effects of the straw returning method and potassium fertilizer dosage on the lodging resistance mechanical properties of spring maize stems

As shown in Table S3, the effects of the single factor, two-factor interactions and three-factor interactions of the straw returning method, potassium fertilizer dosage, and ecological region on the maize stem puncture strength, compressive strength and bending strength were extremely significant.

**Table S3.**
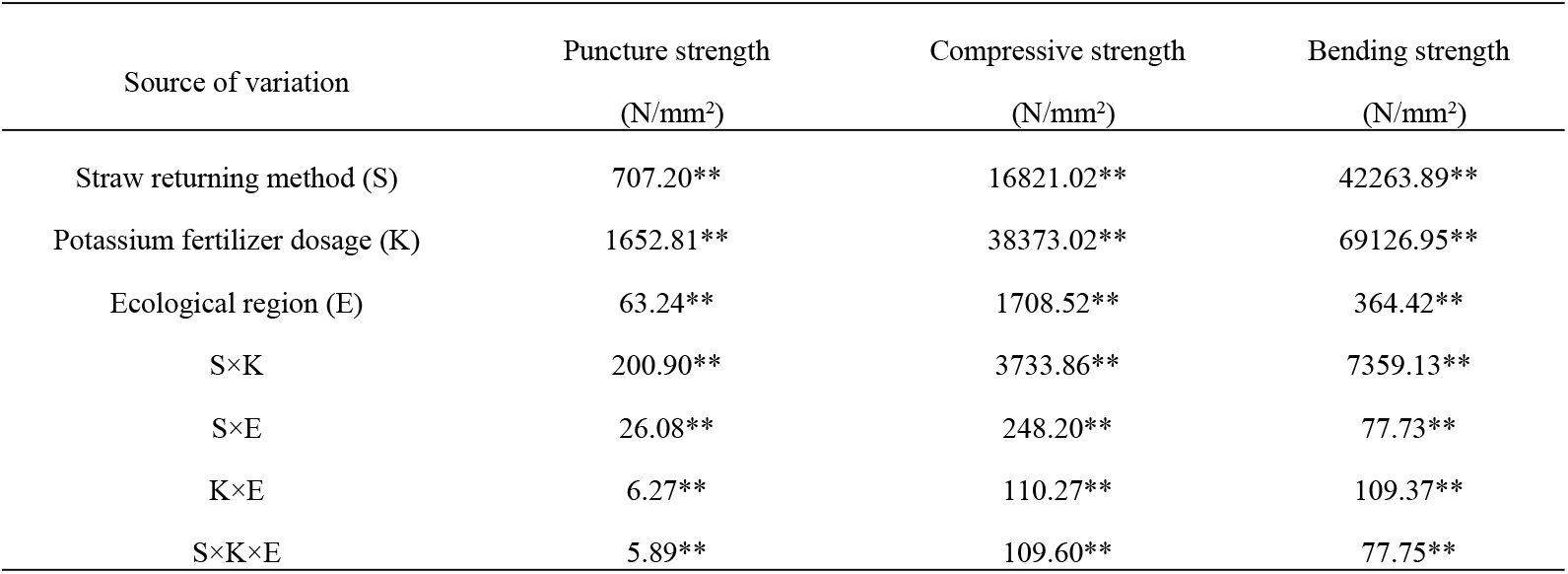
ANOVA results for maize stem lodging resistance mechanical properties under different straw returning methods and potassium fertilizer treatments

As shown in Figure 1 and Table S4, the maize stem puncture strength, stem bending strength and stem compressive strength varied widely between the treatments. When straw returning with no potassium application was compared to straw returning with potassium application in the Tumochuan Plain irrigation area, Hetao Plain irrigation area, and Lingnan warm dry zone, the treatment with potassium increased the stem puncture strength by 22.06%, 26.73%, and 21.17%, respectively, whereas the stem compressive strength was increased by 20.72%, 24.14%, and 20.83%, respectively, and the stem bending strength was increased by 28.82%, 32.56%, and 27.44%, respectively. With no straw returning treatment, the stem puncture strength in the Tumochuan Plain irrigation area, Hetao Plain irrigation area, and Lingnan warm dry zone was increased by 11.51%, 11.91%, and 12.28%, respectively, with potassium application in comparison with no potassium application, whereas the stem compressive strength was increased by 9.99%, 12.37%, and 14.36%, respectively, and the stem bending strength was increased by 16.81%, 16.87%, and 16.55%, respectively. In the Tumochuan Plain irrigation area, straw returning increased the effectiveness of potassium application on the stem puncture strength, stem compressive strength and stem bending strength by 10.55%, 14.82%, and 8.89%, respectively. In the Hetao Plain irrigation area, straw returning increased the effectiveness of potassium application on the stem puncture strength, stem compressive strength and stem bending strength by 10.73%, 11.76%, and 6.47%, respectively. In the Lingnan warm dry zone, straw returning increased the effectiveness of potassium application on the stem puncture strength, stem compressive strength and stem bending strength by 12.01%, 15.69%, and 10.89%, respectively.

**Figure 1.**
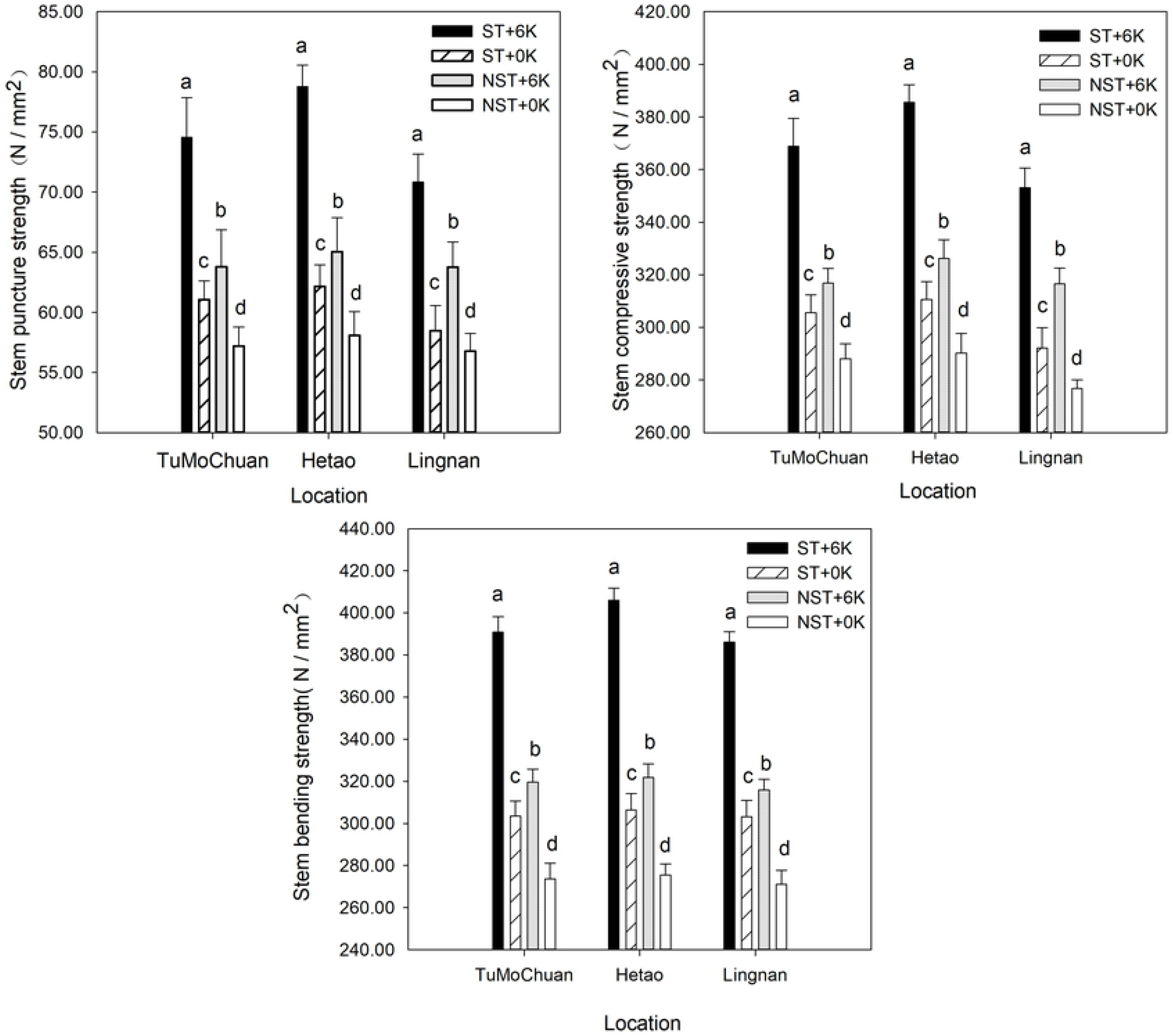
Stem lodging resistance mechanical properties of spring maize under different straw returning methods and potassium fertilizer treatments

**Table S4.** Effects of the interaction of the straw returning method and potassium fertilizer dosage on the lodging resistance mechanical properties of maize stems in different ecological regions

| Ecological region | Straw returning method | Potassium fertilizer dosage | Puncture strength (N/mm<sup>2</sup>) | Compressive strength(N/mm<sup>2</sup>) | Bending strength(N/mm<sup>2</sup>) |
| --- | --- | --- | --- | --- | --- |
| Tumochuan Plain<br>irrigation area | ST | 6K | 74.55±3.31a | 368.84±10.63a | 390.83±7.28a |
|  |  | 0K | 61.06±1.56c | 305.53±6.83c | 303.42±7.24c |
|  | NST | 6K | 63.80±3.06b | 316.83±5.56b | 319.63±6.18b |
|  |  | 0K | 57.21±1.57d | 288.08±5.68d | 273.67±7.40d |
|  | Under ST, the percentage increase of 6K compared to 0K % |  | 22.06 | 20.72 | 28.82 |
|  | Under NST, the percentage increase of 6K compared to 0K % |  | 11.51 | 9.99 | 16.81 |
|  | ST increases the percentage of K effectiveness % |  | 10.55 | 10.73 | 12.01 |
| Hetao Plain<br>irrigation area | ST | 6K | 78.78±1.79a | 385.52±6.70a | 406.06±5.76a |
|  |  | 0K | 62.17±1.79c | 310.59±6.80c | 306.41±7.68c |
|  | NST | 6K | 65.04±2.84b | 326.17±7.18b | 321.84±6.36b |
|  |  | 0K | 58.10±1.97d | 290.28±7.36d | 275.40±5.37d |
|  | Under ST, the percentage increase of 6K compared to 0K % |  | 26.73 | 24.14 | 32.56 |
|  | Under NST, the percentage increase of 6K compared to 0K % |  | 11.91 | 12.37 | 16.87 |
|  | ST increases the percentage of K effectiveness % |  | 14.82 | 11.76 | 15.69 |
| Lingnan warm dry<br>zone | ST | 6K | 70.85±2.31a | 353.05±7.53a | 386.13±4.82a |
|  |  | 0K | 58.48±2.09c | 292.22±7.71c | 303.09±7.89c |
|  | NST | 6K | 63.76±2.09b | 316.62±5.92b | 315.86±5.15b |
|  |  | 0K | 56.78±1.48d | 276.85±3.19d | 271.05±6.60d |
|  | Under ST, the percentage increase of 6K compared to 0K % |  | 21.17 | 20.83 | 27.44 |
|  | Under NST, the percentage increase of 6K compared to 0K % |  | 12.28 | 14.36 | 16.55 |
|  | ST increases the percentage of K effectiveness % |  | 8.89 | 6.47 | 10.89 |

### Effects of the straw returning method and potassium fertilizer dosage on maize grain quality

As shown in Table S5, the effects of variation of the straw returning method, potassium fertilizer dosage, and ecological region on the protein content, starch content, crude fat content and water content of grains all reached an extremely significant level. The effects of the interaction of the straw returning method and potassium fertilizer dosage and the interaction of the potassium fertilizer dosage and ecological region on the indicators mentioned above reached an extremely significant level. The effects of the interaction of the straw returning method and ecological region on the protein content, starch content, and crude fat content of maize grains reached extremely significant levels. The effect of the interaction of the straw returning method, potassium fertilizer dosage and ecological region on the water content of grains was significant.

**Table S5.** ANOVA results for maize grain quality under different straw returning methods and potassium fertilizer treatments

| Source of variation | Protein content of<br>grains (%) | Starch content of<br>grains (%) | Crude fat content of<br>grains (%) | Water content of<br>grains (%) |
| --- | --- | --- | --- | --- |
| Straw returning method (S) | 2.66** | 2.56** | 0.73** | 1.24** |
| Potassium fertilizer dosage (K) | 12.51** | 12.05** | 3.07** | 4.19** |
| Ecological region (E) | 2.65** | 17.78** | 0.74** | 0.89** |
| S×K | 0.68** | 0.25** | 0.06** | 0.55** |
| S×E | 0.08** | 0.09** | 0.01** | 0.01ns |
| K×E | 0.09** | 0.09** | 0.01** | 0.04** |
| S×K×E | 0.02ns | ns | ns | 0.01* |

As shown in Figure 2 and Table S6, the protein content, starch content, crude fat content and water content of maize grains differed significantly among the treatments. In the Tumochuan Plain irrigation area, Hetao Plain irrigation area, and Lingnan warm dry zone, straw returning treatment increased the protein content of grains by 11.78%, 13.68%, and 13.53%, respectively, with potassium application in comparison with no potassium application, whereas the starch content of grains increased by 1.34%, 1.68%, and 1.31%, while the crude fat content of grains increased by 12.71%, 14.72%, and 14.73%, and the water content of grains decreased by 7.30%, 7.22%, and 7.66%. With no straw returning treatment, the protein content of grains in the Tumochuan Plain irrigation area, Hetao Plain irrigation area, and Lingnan warm dry zone increased by 7.52%, 10.19%, and 7.03%, respectively, with potassium application in comparison with no potassium application, whereas the starch content of grains increased by 1.01%, 1.25%, and 0.99%, while the crude fat content of grains increased by 10.21%, 12.62%, and 10.30%, and the water content of grains decreased by 2.63%, 2.67%, and 4.91%. In the Tumochuan Plain irrigation area, Hetao Plain irrigation area, and Lingnan warm dry zone, straw returning increased the effectiveness of potassium application on the protein content of grains by 4.26%, 3.49%, and 6.50%, respectively, whereas its effectiveness on the starch content of grains increased by 0.33%, 0.43%, and 0.32%, while its effectiveness on the crude fat content of grains increased by 2.51%, 2.09%, and 4.43%, and its effectiveness on the water content of grains increased by 4.67%, 4.55%, and 2.75%.

**Figure 2.**
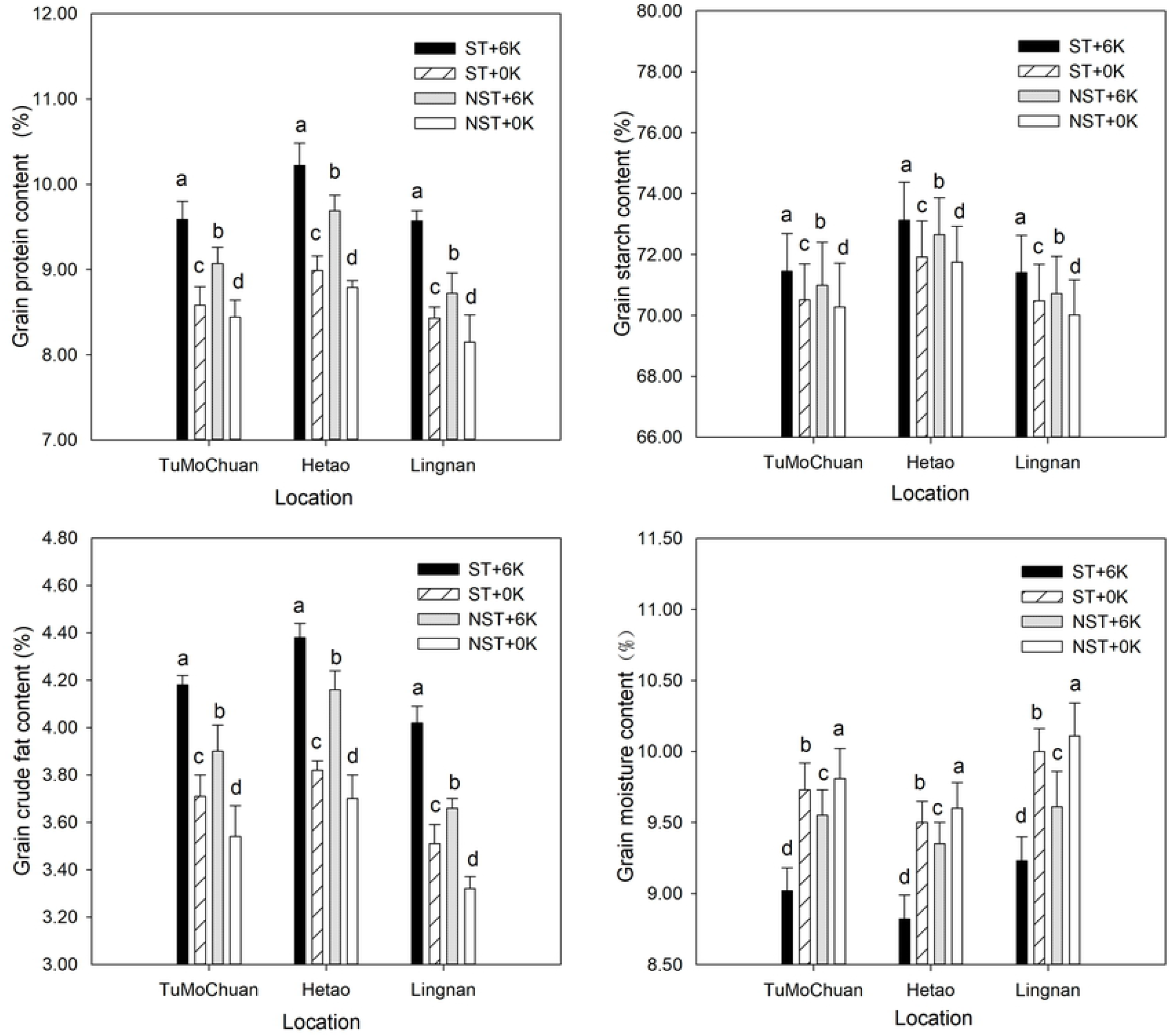
Grain quality of spring maize under different straw returning methods and potassium fertilizer treatments

**Table S6.** Effects of the interaction of the straw returning method and potassium fertilizer dosage on maize grain quality in different ecological regions

| Ecological region | Straw returning method | Potassium fertilizer dosage | Protein content of<br>grains (%) | Starch content of<br>grains (%) | Crude fat content of<br>grains (%) | Water content of<br>grains (%) |
| --- | --- | --- | --- | --- | --- | --- |
| Tumochuan Plain<br>irrigation area | ST | 6K | 9.59±0.21a | 71.45±1.24a | 4.18±0.04a | 9.02±0.16d |
|  |  | 0K | 8.58±0.22c | 70.51±1.18c | 3.71±0.09c | 9.73±0.19b |
|  | NST | 6K | 9.07±0.19b | 70.99±1.41b | 3.90±0.11b | 9.55±0.18c |
|  |  | 0K | 8.44±0.20d | 70.28±1.43d | 3.54±0.13d | 9.81±0.21a |
|  | Under ST, the percentage increase of 6K compared to 0K % |  | 11.78 | 1.34 | 12.71 | 7.30 |
|  | Under NST, the percentage increase of 6K compared to 0K % |  | 7.52 | 1.01 | 10.21 | 2.63 |
|  | ST increases the percentage of K effectiveness % |  | 4.26 | 0.33 | 2.51 | 4.67 |
| Hetao Plain<br>irrigation area | ST | 6K | 10.22±0.26a | 73.13±1.24a | 4.38±0.06a | 8.82±0.17d |
|  |  | 0K | 8.99±0.17c | 71.92±1.18c | 3.82±0.04c | 9.50±0.15b |
|  | NST | 6K | 9.69±0.18b | 72.65±1.22b | 4.16±0.08b | 9.35±0.15c |
|  |  | 0K | 8.79±0.08d | 71.75±1.17d | 3.70±0.10d | 9.60±0.18a |
|  | Under ST, the percentage increase of 6K compared to 0K % |  | 13.68 | 1.68 | 14.72 | 7.22 |
|  | Under NST, the percentage increase of 6K compared to 0K % |  | 10.19 | 1.25 | 12.62 | 2.67 |
|  | ST increases the percentage of K effectiveness % |  | 3.49 | 0.43 | 2.09 | 4.55 |
| Lingnan warm dry<br>zone | ST | 6K | 9.57±0.12a | 71.40±1.22a | 4.02±0.07a | 9.23±0.17d |
|  |  | 0K | 8.43±0.13c | 70.48±1.20c | 3.51±0.08c | 10.00±0.16b |
|  | NST | 6K | 8.72±0.24b | 70.72±1.22b | 3.66±0.04b | 9.61±0.25c |
|  |  | 0K | 8.15±0.32d | 70.02±1.15d | 3.32±0.05d | 10.11±0.23a |
|  | Under ST, the percentage increase of 6K compared to 0K % |  | 13.53 | 1.31 | 14.73 | 7.66 |
|  | Under NST, the percentage increase of 6K compared to 0K % |  | 7.03 | 0.99 | 10.30 | 4.91 |
|  | ST increases the percentage of K effectiveness % |  | 6.50 | 0.32 | 4.43 | 2.75 |

### Effects of straw returning method and potassium fertilizer dosage on maize yield

As shown in Table S7, the effects of the single factor, two-factor interaction and three-factor interactions of the straw returning method, potassium fertilizer dosage, and ecological region on the maize grain number per ear, 1000-grain weight, water content and yield all reached a significant or extremely significant level, whereas their effects on ear number did not reach a significant level.

**Table S7.** ANOVA results for maize yield and yield component factors under different straw returning methods and potassium fertilizer treatments

| Source of variation | Ears per<br>hectare<br>(fringe/hm<sup>2</sup>) | Grains per spike<br>(grain/ fringe) | 1000-seed<br>weight (g) | Water<br>content<br>(%) | Yield<br>(kg/hm<sup>2</sup>) |
| --- | --- | --- | --- | --- | --- |
| Straw returning method (S) | ns | 2485.84** | 7117.71** | 1.54** | 21187480.00** |
| Potassium fertilizer dosage (K) | ns | 9845.77** | 16246.02** | 3.93** | 56653950.00** |
| Ecological region (E) | ns | 1875.02** | 12030.89** | 0.27** | 25155970.00** |
| S×K | ns | 733.60** | 1475.11** | 0.48** | 4589876.00** |
| S×E | ns | 33.74* | 337.56** | 0.03** | 1029908.00** |
| K×E | ns | 60.24** | 113.48** | 0.03** | 644030.00** |
| S×K×E | ns | 28.79ns | 100.11** | 0.02** | 399520.00** |

As shown in Table 5, the maize grain number per ear, 1000-grain weight, water content and yield were significantly different among the treatments. In the Tumochuan Plain irrigation area, Hetao Plain irrigation area, and Lingnan warm dry zone, straw returning with potassium application increased the grain number per ear by 5.43%, 6.56%, and 4.74%, respectively, in comparison with no potassium application. With no straw returning treatment, the grain number per ear increased by 3.27%, 3.33%, and 3.04%, respectively, with potassium application in comparison with no potassium application. Straw returning increased the effectiveness of potassium application on the grain number per ear by 2.15%, 3.22%, and 1.69%, respectively. Under the straw returning treatment, the maize 1000-grain weight in the Tumochuan Plain irrigation area, Hetao Plain irrigation area, and Lingnan warm dry zone increased by 14.60%, 14.79%, and 11.13%, respectively, with potassium application in comparison with no potassium application. With no straw returning, the maize 1000-grain weight increased by 8.18%, 6.82%, and 7.83%, respectively, with potassium application in comparison with no potassium application. Straw returning increased the effectiveness of potassium application on the maize 1000-grain weight by 6.42%, 7.97% and 3.29%, respectively.

**Table 5.** Maize yield and yield component factors under different straw returning methods and potassium fertilizer treatments

| Ecological region | Straw returning method | Potassium fertilizer dosage | Spike<br>(fringes /hm<sup>2</sup>) | Grain number per spike<br>(grains/fringe) | 1000-grain weight (g) | Water content (%) | Yield (kg/hm<sup>2</sup>) | Variation coefficient |
| --- | --- | --- | --- | --- | --- | --- | --- | --- |
| Tumochuan Plain irrigation area | ST | 6K | 79473±814.81a | 615.48±15.29a | 357.90±14.12a | 18.97±0.40d | 14181.86±608.93a | 4.29 |
|  |  | 0K | 79902±453.16a | 583.80±14.00c | 312.36±13.59c | 19.66±0.38b | 11705.04±584.26c | 4.99 |
|  | NST | 6K | 80682±973.64a | 597.12±15.07b | 328.50±11.41b | 19.48±0.37c | 12746.54±665.25b | 5.22 |
|  |  | 0K | 79920±1950.49a | 578.20±14.52d | 303.76±13.01d | 19.76±0.39a | 11263.54±645.34d | 5.73 |
|  | Under ST, the percentage increase of 6K compared to 0K % |  |  | 5.43 | 14.60 | 3.51 | 21.20 | 13.98 |
|  | Under NST, the percentage increase of 6K compared to 0K % |  |  | 3.27 | 8.18 | 1.42 | 13.20 | 8.91 |
|  | ST increases the percentage of K effectiveness % |  |  | 2.15 | 6.42 | 2.09 | 8.00 | 5.07 |
| Hetao Plain irrigation area | ST | 6K | 79842±528.46a | 632.20±13.73a | 392.46±14.70a | 18.75±0.31d | 16091.95±575.74a | 3.58 |
|  |  | 0K | 79359±986.05a | 593.32±14.46c | 342.18±18.76c | 19.56±0.41b | 12949.52±554.12c | 4.28 |
|  | NST | 6K | 79566±671.79a | 606.76±14.87b | 347.30±11.79b | 19.37±0.37c | 13519.84±608.29b | 4.50 |
|  |  | 0K | 78924±751.09a | 587.16±11.61d | 325.28±14.68d | 19.74±0.42a | 12096.89±599.79d | 4.96 |
|  | Under ST, the percentage increase of 6K compared to 0K % |  |  | 6.56 | 14.79 | 4.17 | 24.31 | 16.39 |
|  | Under NST, the percentage increase of 6K compared to 0K % |  |  | 3.33 | 6.82 | 1.89 | 11.81 | 9.26 |
|  | ST increases the percentage of K effectiveness % |  |  | 3.22 | 7.97 | 2.27 | 12.50 | 7.13 |
| Lingnan warm dry zone | ST | 6K | 80256±672.71a | 604.64±15.99a | 326.78±12.21a | 19.17±0.37d | 12815.56±571.99a | 4.46 |
|  |  | 0K | 80313±994.61a | 577.36±17.97c | 294.12±12.48c | 19.74±0.41b | 10945.03±585.23c | 5.35 |
|  | NST | 6K | 79935±1927.75a | 588.84±14.57b | 306.24±12.86b | 19.54±0.43c | 11598.04±660.44b | 5.69 |
|  |  | 0K | 79452±2043.51a | 571.48±15.66d | 284.02±12.44d | 19.89±0.42a | 10333.19±646.47d | 6.26 |
|  | Under ST, the percentage increase of 6K compared to 0K % |  |  | 4.74 | 11.13 | 2.88 | 17.14 | 16.53 |
|  | Under NST, the percentage increase of 6K compared to 0K % |  |  | 3.04 | 7.83 | 1.72 | 12.27 | 8.98 |
|  | ST increases the percentage of K effectiveness % |  |  | 1.69 | 3.29 | 1.15 | 4.87 | 7.55 |

Under the straw returning treatment, the water content of grains in the Tumochuan Plain irrigation area, Hetao Plain irrigation area, and Lingnan warm dry zone was reduced by 3.51%, 4.17%, and 2.88%, respectively, with potassium application in comparison with no potassium application. With no straw returning treatment, the water content of grains was reduced by 1.42%, 1.89%, and 1.72%, respectively, with potassium application in comparison with no potassium application. Straw returning increased the effectiveness of potassium application on the water content of grains by 2.09%, 2.27% and 1.15%, respectively.

Under the straw returning treatment, the maize yield in the Tumochuan Plain irrigation area, Hetao Plain irrigation area, and Lingnan warm dry zone increased by 21.20%, 24.31%, and 17.14%, respectively, with potassium application in comparison with no potassium application. With no straw returning treatment, the maize yield increased by 13.20%, 11.81%, and 12.27%, respectively, with potassium application in comparison with no potassium application. Straw returning increased the effectiveness of potassium application on maize yield by 8.00%, 12.50%, and 4.87%, respectively.

Under the straw returning treatment, the maize yield variation coefficient in the Tumochuan Plain irrigation area, Hetao Plain irrigation area, and Lingnan warm dry zone was reduced by 13.98%, 16.39%, and 16.53%, respectively, with potassium application in comparison with no potassium application. With no straw returning treatment, the maize yield variation coefficient was reduced by 8.91%, 9.26%, and 8.98%, respectively, with potassium application in comparison with no potassium application. Straw returning increased the effectiveness of potassium application on the maize yield variation coefficient by 5.07%, 7.13%, and 7.55%, respectively. Straw returning improved the effectiveness of potassium application on the maize grain number per ear, 1000-grain weight, water content, yield and yield variation coefficient.

## Discussion

### Effects of straw returning combined with potassium fertilizer on the morphological indicators of spring maize stems and stem phenotypic traits

Both straw returning and the application of potassium fertilizer can promote the growth and development of spring maize; the maize plant height, stem diameter, dry matter accumulation of the aerial part of the plant, water content and brix of stems were all increased to varying degrees by these practices [25-27]. In this study, we found that straw returning with potassium fertilizer improved spring maize stem morphological indicators and stem phenotypic traits. The effects of the treatments on each of the tested factors were ranked as follows: ST+6K>NST+6K>ST+0K>NST+0K. These findings are consistent with some studies [28], but not with others [29]. This inconsistency with previously reported results may be due to differences in the time frame of each study. The present study utilized straw returning for consecutive years, while Huijuan Ma conducted experiments with short-term straw returning.

### Effects of straw returning with potassium fertilizer on the mechanical properties of spring maize stems

Both straw returning and potassium fertilizer can significantly enhance the puncture strength, compressive strength and bending strength of spring maize stems, thus increasing maize lodging resistance [30,31]. In this study, straw returning combined with potassium fertilizer significantly improved the mechanical properties of spring maize stems, with the effectiveness of the treatments ranked as follows: ST+6K>NST+6K>ST+0K>NST+0K. These findings are consistent with previous studies [32]. Potassium fertilizer promotes the absorption of potassium by maize, and the absorbed potassium is primarily distributed in the stems, where it contributes to the flexural resistance of stems and has the potential to supplement soil potassium via straw returning at a later stage. Straw returning can increase the available potassium content of the surface soil, yet its direct effect on the aerial part of maize plants is not as pronounced as that of potash application.

### Effects of straw returning combined with potassium fertilizer on the grain quality of spring maize

Both straw returning and potassium fertilizer can improve the protein content, crude fat content and starch content of maize grains, and thus enhance maize grain quality [33,34]. In this study, straw returning combined with potassium fertilizer improved spring maize grain quality, with the effectiveness of the treatments ranked as follows: ST+6K>NST+6K>ST+0K>NST+0K.

### Effects of straw returning with potassium fertilizer on spring maize yield

Crop yield is a primary parameter used in the evaluation of the effects of fertilizer application on soil productivity [35]. Potassium fertilizer, straw returning and the combination of straw returning with potassium fertilizer have all been shown to significantly increase spring maize yield [36-38]. In this study, straw returning with potassium fertilizer increased spring maize yield, with the effectiveness of the treatments ranked as follows: ST-6K>NST-6K>ST-0K>NST-0K. These findings are consistent with previous studies [39,40]. When potassium fertilizer enters the soil, it is converted into soil-available potassium, which can be directly absorbed and used by maize. In comparison with potassium contained in potassium fertilizer, potassium in maize straw is more easily fixed by the soil and is not readily released. Straw enters the soil after it is returned to the field, and potassium in returned straw must be released by the action of soil microorganisms and enzymes via a long and complex decomposition process. Therefore, potassium in returned straw cannot satisfy the potassium required for the growth and development of maize in the current season. Thus, in comparison with direct application of potassium fertilizer, short-term straw returning has a weaker effect on maize yield.

The variation coefficient of repeated fluctuations in crop yield is an important indicator used to evaluate the disadvantages and advantages of fertilization systems. The main factors affecting the maize yield variation coefficient are soil fertility and soil basic productivity. When the variation coefficient is relatively small, stability is relatively high [41]. In this experiment, the selected optimal model for stable yield was straw returning with potassium fertilizer, and the spring maize yield variation coefficients of the treatments were ranked as follows: ST-6K<ST-0K<NST-6K<NST-0K. These findings are consistent with previous studies [42]. In this study, stable yield was achieved primarily because straw returning with potassium fertilizer improved the soil structure and physicochemical properties, increased soil nutrient levels and soil basic productivity, and provided a suitable environment for the growth and development of spring maize, thus reducing the maize yield variation coefficient and increasing yield stability.

The spring maize stem morphological indicators, stem phenotypic traits, stem mechanical properties, grain quality, and yield varied among the three tested ecological regions in Inner Mongolia, yet the trends in the changes in relevant measurements in each of these areas due to the application of straw returning with potassium fertilizer were extremely similar. Among the tested ecological regions, each of these factors was superior in the Hetao Plain irrigation area, followed by the Tumochuan Plain irrigation area, and finally the Lingnan warm dry zone. These differences were mainly due to differences in factors such as basic soil productivity and climatic conditions (sunshine hours, temperature, and rainfall) among the regions.

## Conclusions

Among the different treatments, straw returning with potassium fertilizer demonstrated the best effect. Straw returning significantly improved the effectiveness of potassium application on the morphological indicators of spring maize stems, stem phenotypic traits, stem mechanical properties, grain quality, and yield. In the Tumochuan Plain irrigation area, Hetao Plain irrigation area, and Lingnan warm dry zone, straw returning increased the stem dry weight, water content of stems, brix of stems, stem puncture strength, stem compressive strength, stem bending strength, protein content of grains, starch content of grains, crude fat content of grains, water content of grains, yield and yield variation coefficient by 8.89%-11.65%, 1.09%-1.55%, 4.35%-4.50%, 8.89%-14.82%, 6.47%-11.76%, 10.89%-15.69%, 3.49%-6.50%, 0.32%-0.43%, 2.09%-4.43%, 2.75%-4.67%, 4.87%-12.50% and 5.07%-7.55%, respectively. Straw returning with potassium fertilizer improved spring maize stem lodging resistance, while improving grain quality and achieving stable and high yields. This study provides a theoretical basis for high-yield cultivation of stress-resistant spring maize in Inner Mongolia.

## Acknowledgments

We would like to thank the Maize High-Yield and High-Efficiency Cultivation Team for field and data collection.

## Author Contributions

Funding acquisition: Julin Gao, Jiying Sun.

Investigation: Yafang Fan.

Methodology: Jiying Sun, Jian liu, Zhijun Su, Shuping Hu, Zhigang Wang, Xiaofang Yu.

Conceptualization: Julin Gao, Jiying Sun, Jian liu, Yafang Fan.

Writing – original draft: Yafang Fan, Jian liu.

Writing – review & editing: Julin Gao, Jiying Sun.

